# Ethanol decreases *Pseudomonas aeruginosa* flagellar motility through a cyclic-di-GMP- and stator-dependent pathway

**DOI:** 10.1101/430884

**Authors:** Kimberley A. Lewis, Amy E. Baker, Annie I. Chen, Colleen E. Harty, Sherry L. Kuchma, George A. O’Toole, Deborah A. Hogan

## Abstract

*Pseudomonas aeruginosa* frequently encounters microbes that produce bioactive metabolites including ethanol. At concentrations that do not affect growth, we found that ethanol reduces *P. aeruginosa* motility by 30% in a swim agar assay and this decrease is accompanied by a 2.5-fold increase in levels of cyclic diguanylate (c-di-GMP), a second messenger that represses motility, in planktonic cells. A screen of mutants lacking genes involved in c-di-GMP metabolism identified SadC and GcbA as diguanylate cyclases involved in swim zone reduction by ethanol and ethanol-induced c-di-GMP production. The reduction of swimming in response to ethanol also required the stator set, MotAB, two PilZ-domain proteins (FlgZ and PilZ), PilY1-a proposed surface-sensing protein, and PilMNOP, which comprises the pilus alignment complex and these proteins have been previously implicated in the control of motility on agar surfaces. Microscopic analysis of the fraction of quiescent cells in swim medium showed that ethanol decreased the portion of motile cells in the wild type, but had opposite effects in the ∆*pilY1*, ∆*pilMNOP*, ∆*motAB*, and ∆*pilZ*∆*flgZ* mutants. Together, these data indicate ethanol induces a regulated change in motility in planktonic cells at concentrations similar to those produced by other microbes. We propose that this ethanol-responsiveness may contribute to the co-localization of *P. aeruginosa* with ethanol-producing microbes.

## Importance

Ethanol is an important, biologically active molecule produced by many bacteria and fungi. It has also been identified as a potential marker for disease state in cystic fibrosis. In line with previous data that show that ethanol promotes biofilm formation by *Pseudomonas aeruginosa*, here we report that ethanol also induces cyclic-di-GMP levels in planktonic cells and reduces swimming motility using some of the same proteins involved in surface sensing. We propose that these data may provide insight into how microbes can influence *P. aeruginosa* localization and surface association in the context of infection and in other polymicrobial settings.

## Introduction

Ethanol, in addition to being a common fermentation product and a carbon source, can also serve as a signaling molecule for many microbes. For example, fungal gardens formed as part of a symbiosis between ambrosia beetles and their fungal symbionts, *Ambrosiella* and *Raffaelea*, are preferentially localized to sites with higher ethanol (1). In the parasite *Toxoplasma gondii*, low concentrations of ethanol (<200 mM or 1.2%) facilitates an increase in the second messenger inositol 1, 4, 5-triphosphate, resulting in increased intracellular calcium and increased host colonization (2, 3). In *Acinetobacter baumannii*, a Gram-negative opportunistic pathogen, ethanol causes an increase in virulence and an increase in carbohydrate leading to biofilm formation and repression of motility through mechanisms not yet described (4). In *Pseudomonas aeruginosa*, exogenous ethanol and ethanol produced by the fungus *Candida albicans* alters phenazine production and promotes biofilm formation on plastic and airway cells (5). Ethanol leads to increased Pel matrix production and decreased surface motility, two factors that are necessary for biofilm formation and maturation (5–7).

The many responses to ethanol are not surprising considering its frequent production as a fermentation product by many bacteria and fungi, including in host infection settings. One example where microbial ethanol is detected is in the polymicrobial lung infections associated with cystic fibrosis (CF), a genetic disorder that results in an accumulation of thick mucus in the airways (8–10). In addition to *P. aeruginosa*, CF lung infections often contain other microbes, many of which are capable of producing ethanol (11). Metabolomic and NMR studies examining the bronchoalveolar lavage fluid (BALF) and exhaled breath condensate (EBC) of patients with CF indicate that volatiles such as ethanol are present in the CF lung at varying amounts depending on the state of disease (stable vs. during exacerbation), and therefore may serve as biomarkers of disease (8, 9, 12, 13).

Previous studies found that in surface-associated *P. aeruginosa*, ethanol stimulates an increase in the level of the second messenger, cyclic-di-GMP (c-di-GMP), in part through WspR, a diguanylate cyclase (DGC) (5). WspR is among the forty enzymes in *P. aeruginosa* thought to metabolize c-di-GMP, including other DGCs (14), c-di-GMP-degrading phosphodiesterases (PDEs) (15–17), and proteins that possess both activities. In *P. aeruginosa* and other pseudomonads, c-di-GMP metabolic enzymes have additional domains (e.g. PAS, REC, HAMP, CAHCE and GAF) that can sense external stimuli or promote protein-protein interactions in order to modulate enzyme activities at appropriate times (18–20), or in response to specific cues (19). In addition to the c-di-GMP metabolic enzymes, *P. aeruginosa* has thirteen effector proteins that bind c-di-GMP at various affinities to affect many behaviors including biofilm formation and motility (21, 22).

One of the roles of high c-di-GMP in *P. aeruginosa* is the down-regulation of flagellar motility (5, 7, 22–26). In pseudomonads and other bacteria, motility repression occurs in multiple ways, including (i) obstruction of the flagellum by exopolysaccharides (27, 28), (ii) transcriptional down-regulation of flagellar gene expression (26, 29), (iii) loss of flagellar rotation by c-di-GMP-bound effector proteins and their interactions with flagellum motor components (26, 30, 31), (iv) sequestration of flagellar motor proteins by c-di-GMP-bound effectors (22, 24), and (v) inhibition of flagellar rotation switching (clockwise vs. counterclockwise) (25, 32, 33). In Gram-negative bacteria, the flagellar motor is composed of two structures, the rotor (FliG, FliM, and FliN), which determines clockwise or counterclockwise rotation (the switch complex), and the stator (MotA and MotB), which generates torque for flagellar rotation powered by proton motive force (34–36). Pseudomonads have a second stator set (MotCD) that is incorporated into the stator complex to facilitate optimal motor function (22, 24, 37, 38). In *P. aeruginosa* the two stator sets have distinct roles: MotAB is required to reduce swarming motility when c-di-GMP levels are high, while MotCD is critical for promoting swimming and swarming motility (24, 37).

In the present study, we outline a pathway by which 1% ethanol represses motility in planktonic *P. aeruginosa* cells. Decreased swimming motility in cells exposed to ethanol is accompanied by a sustained increase in global cellular c-di-GMP pools. Genetic screens found two DGCs, SadC and GcbA, two PilZ-domain effector proteins (FlgZ and PilZ), and the MotAB stator set as components required for ethanol-dependent motility repression. In addition, PilY1, and the PilMNOP proteins, components of the type 4 pili (T4P) machinery that are involved in surface sensing, were also required for the ethanol response in the swim agar assay. Ethanol decreased the portion of motile cells in the wild type by microscopic analysis, but mutants blind to the effects of ethanol (∆*pilZ*∆*flgZ*, ∆*motAB*, ∆*pilY1*, and ∆*pilMNOP*) had an opposite response, meaning cells were more motile in ethanol. Taken together with our previous studies (5), we propose that ethanol, a common metabolite produced by microbes, acts as a signal to rapidly repress *P. aeruginosa* swimming motility in planktonic cells, and thus potentiate biofilm initiation.

## Materials and Methods

### Strains and Media

Strains and plasmids used in this study are listed in Table S3. *P. aeruginosa* PA14 and *E.coli* strains were routinely cultured on lysogeny broth (LB) solidified with 1.5% agar, or in LB broth at 37°C with shaking. Gentamicin (Gm) was used at 60 µg/ml and carbenicillin (Cb) at 700 µg/ml for *P. aeruginosa*. Gm was used at 10 µg/mL for *E. coli*. For *P. aeruginosa* phenotypic assays, either M63 (22 mM KH_2_PO_4_, 40 mM K_2_HPO_4_, and 15 mM (NH_4_)_2_SO_4_) or M8 (42 mM Na_2_PO_4_, 22 mM KH_2_PO_4_, and 8.5 mM NaCl) minimal salts medium supplemented with MgSO_4_ (1 mM), glucose (0.2%), and casamino acids (CAA; 0.5%), as indicated. When stated, 1% (v/v) ethanol (200-proof) was added to cooled medium (~50°C) and equivalent volume of water was added to control cultures. For expression plasmids harboring pBAD promoter, arabinose was added to the culture as needed (0.02 or 0.05%).

### Growth curve of *P. aeruginosa* PA14 wild type in the presence of ethanol

Growth curve analysis was performed by diluting *P. aeruginosa* to an OD_600_ of ~0.01 in six ml M63 medium without and with 1% (v/v) ethanol and incubation at 37°C on a roller drum. OD_600_ was measured at specified time points using a Spectronic 20 spectrophotometer. Each sample type was analyzed in triplicate.

### Molecular techniques

Plasmids were made using previously described homologous recombination in *Saccharomyces cerevisiae* (39). Plasmids were then extracted from the yeast using the ‘smash and grab’ method and electroporated into *E. coli* S-17 cells and confirmed via colony PCR. *E. coli* with confirmed constructs were then conjugated with the indicated *P. aeruginosa* strain to generate in-frame deletion mutants using allelic replacement as previously described (39). Exconjugants were selected on solid LB using gentamycin and nalidixic acid followed by counterselection on 5% sucrose. PCR amplification and DNA sequencing, using primers that flanked the site of deletion, were used to confirm all resulting mutants.

For arabinose-inducible complementation, the gene being complemented was expressed on either pMQ80 (60 µg/ml gentamycin) or pDPM73 (700 µg/ml carbenicillin) plasmid backbones. Confirmed constructs were electroporated into the indicated *P. aeruginosa* strains, selecting for the appropriate antibiotic resistance marker. Arabinose (0.02 or 0.05%) was added to the medium and complementation was confirmed via the indicated phenotypic assay.

### Swimming motility assays

Swim assays were performed as previously described (18). Briefly, M63 medium without and with 1% (v/v) ethanol and solidified with 0.3% agar (swim agar) was poured into petri plates and allowed to dry at room temperature (~25°C) for ~4 h prior to inoculation. Sterile tooth picks were used to inoculate bacteria into the center of the agar without touching the bottom of the plate; liquid cultures grown for 8–16 h were used as inoculum. No more than four strains were assayed per plate. Plates were incubated upright at 37°C in stacks of no more than four plates per stack for 16 h; the swim zone diameter was then measured. *P. aeruginosa* wild type was included in each experiment so that mutant phenotypes could be assessed despite slight day-to-day variation in swim zone diameter. Each strain was inoculated in four replicates and replicate values were averaged to obtain a final swim zone diameter for each strain. All strains were assessed on at least three separate days.

### Twitching motility assays

Twitching motility assays were performed with T-agar medium (10g tryptone, 5g NaCl, and 15g agar in 1L) without and with 1% ethanol in petri plates that were allowed to dry at room temperature for 24 h prior to inoculation. Sterile toothpicks were used to inoculate into the agar until the toothpick touched the bottom of the petri plate; liquid cultures grown for 16 h were used as inoculum. No more than four strains were analyzed per plate and six replicate plates were included in each experiment. Plates were incubated in inverted stacks of four at 37°C for 40 h. To visualize the twitch zone, a spatula was used to gently ease the agar out of the petri plates and two mL of 0.1% (w/v) crystal violet in water was added to each plate and allowed to stand for 10 min. The crystal violet was removed and the plates rinsed with water and allowed to air dry. Twitch zone diameter was measure and recorded. All strains were assessed on at least three separate days.

### Swarming motility assays

Swarm assays were performed as previously described (18). Briefly, M8 medium, without and with 1% ethanol, and with 0.5% agar (swarm agar) was poured into 60 × 15 mm plates and allowed to dry at room temperature for ~4 h prior to inoculation. Each plate was inoculated with 0.5 µL of a liquid culture that was grown for 8–16 h, and the plates incubated face-up at 37°C in stacks of no more than four for 16 h. Each strain was inoculated in four replicates and was assessed on at least three separate days. Images were captured using a Canon EOS Rebel T6i camera and images measured for ethanol-dependent swarm repression.

### Reversal rate measurements

To measure the frequency at which a motile cell changes its direction, we used a modified version of a method that was previously described (25, 40). Briefly, overnight liquid cultures were subcultured 1:100 in five mL M63 medium and incubated at 37°C for 2 h. Once cultures reached exponential phase, they were then diluted 1:1000 in fresh M63 medium and Ficoll was added to a final concentration of 3% to obtain higher viscosity conditions that slowed the swimming cells sufficiently to allow the monitoring of reversal rates and mimic swimming in soft agar. Cells were then exposed to either control medium or medium containing 1% (v/v) ethanol for 15 min. Two hundred and fifty microliter of treated cell culture was next gently pipetted into a 35 mm glass bottom MatTek dish and a glass cover slip was placed over the added culture. Four time lapse movies per strain and condition were captured with dark field using the Nikon Eclipse Ti microscope (Nikon Instruments Inc., Melville, NY) equipped with a 10X objective, a Hamamatsu ORCA-Flash 4.0 camera and Nikon NIS Elements AR 4.13.04 64 bit software. Time lapse movies were 8 s in duration with images captured at 20–25 ms intervals with RAM capture and 50 fps. Fiji ImageJ TrackMate (41) was used to process, analyze and quantify the reversal rates of 40 cells per movie. Movies were advanced frame by frame and individual cells were evaluated for the number of times they changed direction within the field of view and reversal rates were normalized and recorded as reversals per 10 s.

### Microscopic agar motility assay

The population percentages of motile and immobile cells were calculated in 0.3% swim agar. Overnight liquid cultures were subcultured 1:100 in five mL M63 medium and incubated at 37°C for 2 h. Once cultures reached exponential phase, they were then diluted by 1:1000 into freshly-prepared M63 swim agar (0.3%) without and with 1% (v/v) ethanol cooled to ~45°C. Two hundred and fifty microliters of each agar mixture were pipetted into a chamber slide (see Fig. 6A) and allowed to solidify and acclimate to treatment for 30 min. Three to four time lapse movies per chamber slide, with two chamber slides per condition, were captured using the 40X objective on the Nikon Eclipse Ti microscope (Nikon Instruments Inc., Melville, NY) equipped with a Hamamatsu ORCA-Flash 4.0 camera and Nikon NIS Elements AR 4.13.04 64 bit software. Time lapse movies were 8 s in duration with images captured at five ms intervals with RAM capture and 100 fps. Fiji ImageJ TrackMate (41) was used to process, analyze and quantify the percentage of cells that were motile and immobile for the entire duration of each movie. Movies were advanced frame by frame and individual cells were evaluated for movement. All strains were assessed on at least two separate days.

### *In vivo* cyclic-di-GMP quantification

C-di-GMP was measured as previously described (5) with modifications. Overnight liquid cultures were diluted 1:1000 in six mL M63 medium with either 1% ethanol or an equivalent volume of water and grown at 37°C for 16 h on a roller drum. Cultures were then adjusted to similar densities (OD_600_) if necessary. Five mL of each culture was pelleted at 4,500 x g for 15 min at 4°C. Cyclic-di-GMP was extracted by vigorously suspending the pellet in 250 µL of ice cold extraction buffer (40:40:20 MeOH/Acetonitrile/dH_2_O and 0.1 N Formic acid, stored at −20°C) and incubating at −20°C for 1 h with tubes positioned upright. Tubes were then centrifuged briefly prior to transfer of the entirety of each extraction mix to a pre-weighed ice cold 1.5 mL Eppendorf tube. Cell debris was pelleted at 15,682 x g for five min at 4°C, 200 µL of the extracted nucleotide was recovered into a clean 1.5 mL ice cold Eppendorf tube, and samples were each neutralized with 4 µL of 15% NH_4_HCO_3_ per 100 µL of sample. Pellets were dried on high for 1 h and the liquid samples on low overnight using the Savant Speed Vac SC110. The pellet weights were measured to get sample dry weight and the dried liquid samples containing the extracted nucleotides were each suspended in 200 µL HPLC-grade water. Two hundred microliters of each sample was sent to RTSF Mass Spectrometry and Metabolomics Core at Michigan State University for LC-MS-MS analysis. Each strain and treatment condition was analyzed in five replicates.

### Statistical analysis

Unpaired Student *t* test, Two-way ANOVA with multiple comparisons, and One-way ANOVA with multiple comparisons were performed pairwise between the wild type and each strain, as well as ethanol and control conditions, using the GraphPad Prism 6 software (GraphPad, La Jolla, CA).

## Results

### Ethanol represses *P. aeruginosa* PA14 swimming motility independently of catabolism and without reducing growth rate

Exogenous ethanol stimulates *P. aeruginosa* biofilm behaviors, including attachment to glass and plastic, pellicle formation, and microcolony formation on airway cells, in part through stimulation of Pel extracellular matrix production (5, 42). Although *P. aeruginosa* can catabolize ethanol (43, 44), ethanol catabolism is not required for these phenotypes (5) indicating that the ethanol was acting as a signal or stimulus that modulates *P. aeruginosa* biofilm-related behaviors.

To further characterize the response to non-inhibitory concentrations of ethanol, we assessed ethanol effects on flagellar motility using a swim agar assay. We observed that *P. aeruginosa* strain PA14 wild type had a 33% smaller swim zone diameter in the presence of 1% ethanol when compared to the control cultures (49 mm versus 33 mm; p<0.0001) (Fig. 1A). A Δ*flgK* mutant that lacks a flagellum is non-motile and served as a reference strain (Fig. 1A). This reduction in swim zone diameter was not a result of differences in growth as *P. aeruginosa* strain PA14 wild type had similar growth rates in this medium in the absence or presence of 1% ethanol (Fig. S1A). Furthermore, the reduction in swim zone diameter also occurred independently of ethanol catabolism as a Δ*exaA* mutant, which cannot grow with ethanol as a carbon source (5), still showed motility repression when ethanol was added to the medium (Fig. S1B). These data indicated that the ethanol-dependent motility repression observed was not a result of ethanol metabolism or a change in the rate of growth.

**FIG. 1.**
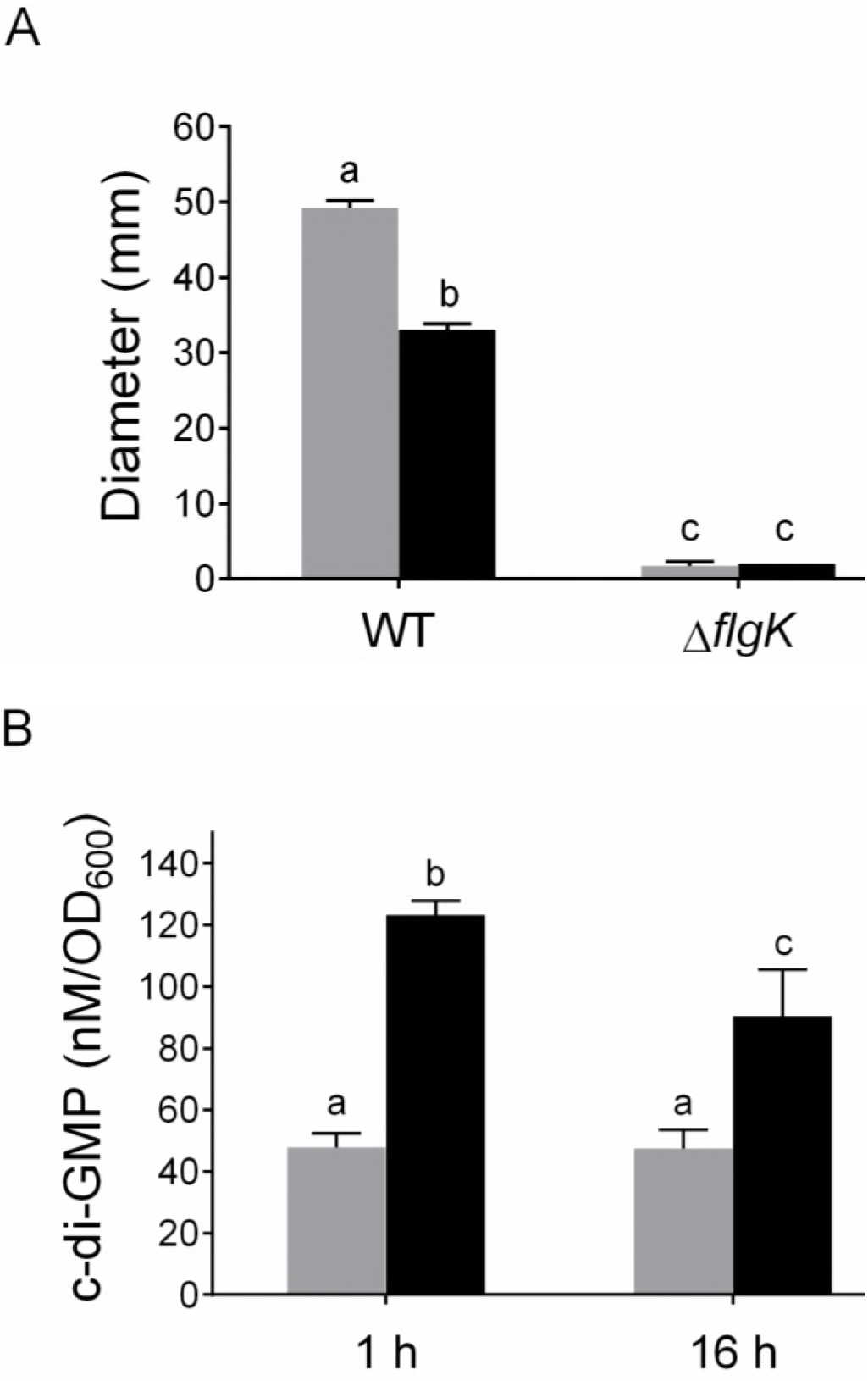
Ethanol decreases swim zone diameter and elicits an early and sustained increase in c-di-GMP. (A) Swim zone diameter of *P. aeruginosa* PA14 wild type and Δ*flgK* in M63 with 0.3% agar (swim agar) without (grey) and with (black) 1% ethanol after 16 h. Bars depict the average swim zone diameter (n=4 replicates). Error bars indicate standard deviations. (B) Quantification of c-di-GMP levels in *P. aeruginosa* PA14 grown in liquid M63 after 1 h and 16 h exposure to 1% ethanol (black) or medium with no ethanol (grey). Bars depict the average of normalized values (n=5 replicates). Error bars indicate standard deviations. Same letters are not significantly different and different letters are significantly different (*P*<0.05) as determined by Two-way ANOVA with multiple comparisons.

### Ethanol elicits an increase in c-di-GMP levels

C-di-GMP is an intracellular signaling molecule that modulates motility (22, 24, 26). When c-di-GMP levels are high, motility is reduced via multiple mechanisms (6, 21, 22, 45–47). In light of the observed decrease in swimming motility (Fig. 1A), we examined the effects of ethanol on c-di-GMP levels in planktonic cells after 1 h and 16 h of growth in medium with 1% ethanol. We found that ethanol caused a 2.6- and 1.9-fold increase (p<0.0001) in c-di-GMP at 1 h and 16 h, respectively (Fig. 1B).

### Ethanol-dependent motility repression is not due to increased Pel and alginate matrix production

Increased c-di-GMP signals have been associated with an increase in alginate and Pel matrix production in *P. aeruginosa* (6, 7). Although ethanol activates WspR-dependent production of Pel polysaccharide matrix (5), neither WspR nor PelA was required for the reduction in swim zone diameter in the presence of ethanol, with a 36.4% and 31.6% (p<0.0001) decrease in their swim zones, respectively (Figs. S2A-B). We did note that the Δ*wspR* mutant had a slightly larger swim zone diameter in control conditions (Fig. S2B). Alginate was also not required for a reduction in swim zone diameter in the presence of ethanol, as two mutants defective in alginate production, Δ*algD* and Δ*algU*, also had similar levels of swim zone reduction (26.7% and 32.6% decrease (p<0.0001), respectively, as the wild type (Fig. S3). These data indicate ethanol-dependent motility repression occurred independently of matrix production.

### A screen of proteins that contribute to c-di-GMP metabolism reveal multiple enzymes involved in the ethanol response

While WspR was found to be required for increased c-di-GMP in response to ethanol in surface-associated cells (5), the Δ*wspR* mutant still showed increased c-di-GMP in planktonic cells in medium with ethanol compared to control medium (Fig. S2C). This suggests that other enzymes are involved in the response to ethanol in planktonic cells. Thus, we screened the collection of the reported *P. aeruginosa* PA14 in-frame deletion mutant library containing mutants lacking each of the 40 known c-di-GMP metabolizing enzymes (18) to identify the gene(s) involved in the ethanol-dependent motility repression. Our primary focus was on mutants that (i) had a swim zone greater than or equal to that of the wild type in control conditions, and (ii) had showed less of a reduction in swim zone diameter when ethanol was present in the medium. Using these criteria, analysis of the data from three independent screens of the mutant collection identified SadC and GcbA as the most promising candidates (Table. S1); data for mutants with swim zone sizes smaller than wild type under control conditions are provided (Table S2), but not pursued as part of these studies. The differences in the magnitude of the effect of ethanol on swim zone size in the Δ*gcbA* and Δ*sadC* single mutants compared to the wild type were small (Fig. 2A and Fig. S4A-B), but could be complemented with the wild-type *gcbA* and *sadC* genes, respectively, in *trans* (Fig. S4A-B). Both SadC and GcbA have been reported to impact c-di-GMP levels (7, 14, 25). Deletion of *wspR* in combination with either *sadC* or *gcbA* did not enhance the resistance of the effects of ethanol on motility (Fig. 2A).

**FIG. 2.**
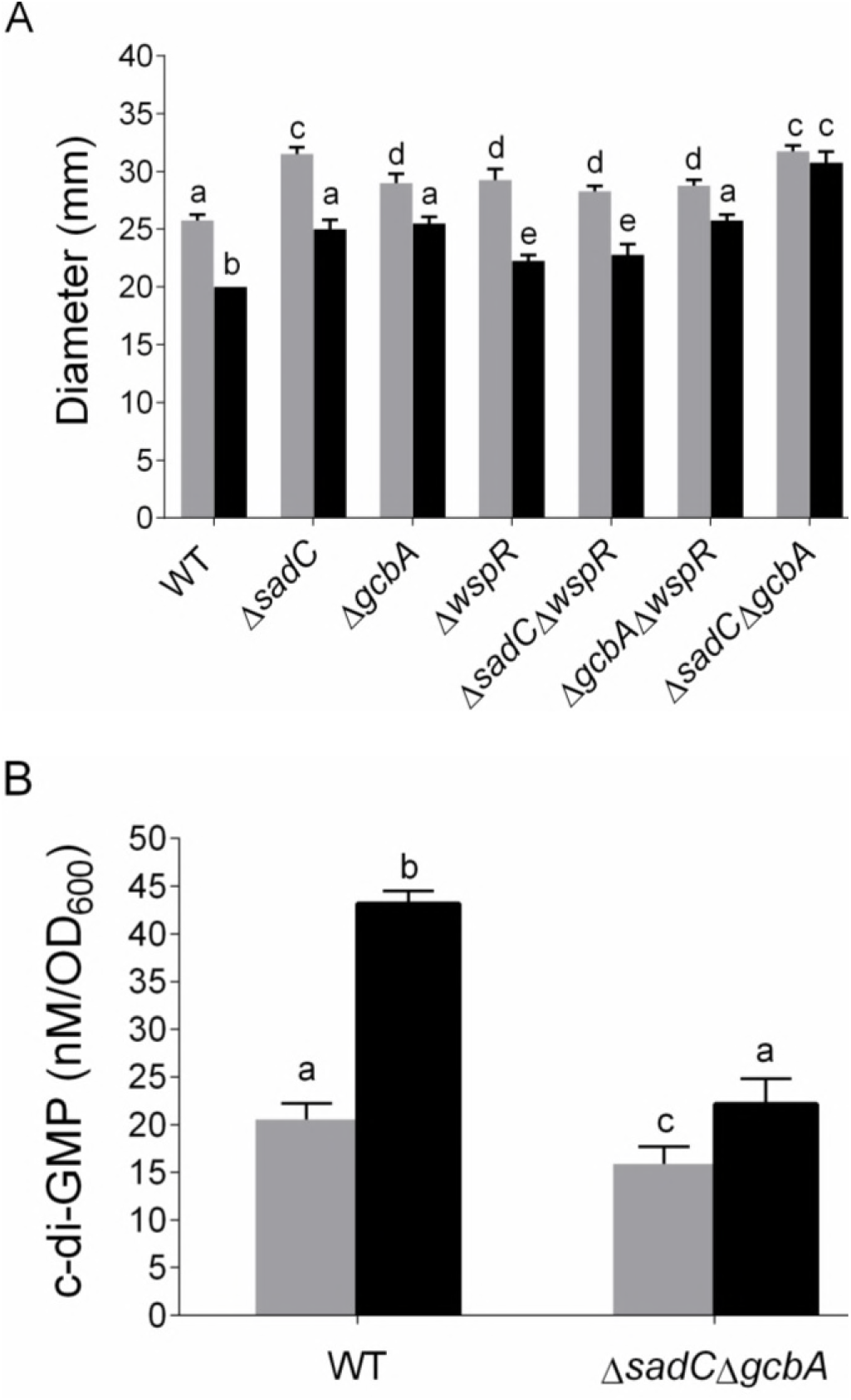
Ethanol effects on motility require two diguanylate cyclases, SadC and GcbA. (A) Swim zone diameter of *P. aeruginosa* PA14 wild type, Δ*sadC*, Δ*gcbA*, Δ*wspR*, Δ*sadC*Δ*wspR*, Δ*gcbA*Δ*wspR*, and Δ*sadC*Δ*gcbA* in swim agar without (grey) and with (black) 1% ethanol measured at 16 h. Error bars indicate standard deviations, n=4 replicates. (B) Quantification of c-di-GMP levels in *P. aeruginosa* PA14 wild type and Δ*sadC*Δ*gcbA*, grown in liquid M63 medium without (grey) and with (black) 1% ethanol for 16 h. Error bars indicate standard deviations, n=5 replicates. Common letters indicate no significant differences, different letters indicate significant differences (*P*-value <0.05) as determined by Multiple t-test corrected using the Holm-Sidak method (A) or One-way ANOVA with multiple comparisons (B).

The effects of SadC and GcbA on changes in motility in response to ethanol were additive as the Δ*sadC*Δ*gcbA* double mutant showed no significant difference in swim zone diameter between medium without and with ethanol (Fig. 2A). The Δ*sadC*Δ*gcbA* mutant also had lower levels of c-di-GMP in planktonic cultures, both in the absence and presence of ethanol, when compared to wild type in the same conditions (Fig. 2B). While the wild type showed 2.1-fold higher levels in c-di-GMP in ethanol-grown cells, the Δ*sadC*Δ*gcbA* mutant only showed a 1.4-fold difference. The small but significant increase that remained in the Δ*sadC*Δ*gcbA* mutant upon growth with ethanol suggests that other enzymes may contribute to changes in cellular c-di-GMP pools when ethanol is present.

### Ethanol induced motility repression requires two PilZ-domain proteins, FlgZ and PilZ

Among the c-di-GMP binding effectors in *P. aeruginosa* are PilZ-domain proteins (47, 48). There are eight known PilZ-domain proteins in *P. aeruginosa*, and some of these proteins have been shown to mediate changes in motility and/or biofilm formation (22, 48). Given that ethanol stimulates c-di-GMP production and motility regulation, we assessed whether one or more of these PilZ-domain proteins might be involved in ethanol-dependent motility repression.

In the absence of ethanol, all eight mutants and the wild type had swim zone diameters that were similar (Fig. 3A). While six of the mutants phenocopied the wild type, two mutants displayed significantly greater swimming motility than that observed for the wild-type strain in the presence of ethanol (Δ*flgZ* and Δ*pilZ*; Fig. 3A). A Δ*flgZ*Δ*pilZ* double mutant had the same level of motility in the presence and absence of ethanol, and thus did not show ethanol-dependent motility repression (Fig. 3B). Interestingly, both PilZ and FlgZ were shown previously to be involved in the repression of swarming motility on agar surfaces in a *P. aeruginosa* strain that had high levels of c-di-GMP due to the absence of a phosphodiesterase, and to regulate flagellar motility in other species (22, 27, 49, 50). Together, these data indicated that PilZ and FlgZ play partially redundant roles in ethanol-dependent motility repression.

**FIG. 3.**
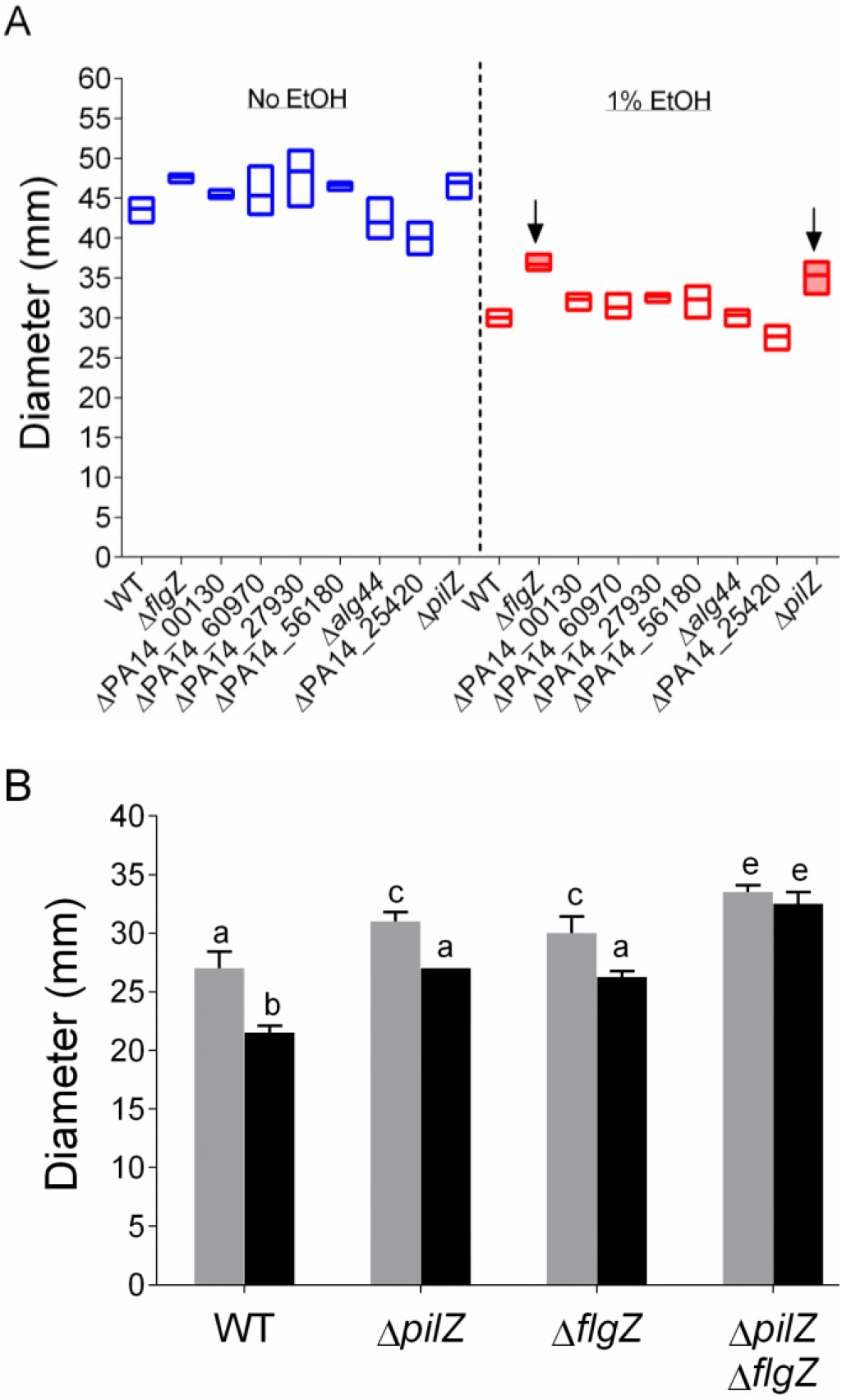
PilZ domain proteins, PilZ and FlgZ, are required for ethanol effects on swim zone diameter. (A) Swim zone diameter for *P. aeruginosa* PA14 wild type, Δ*flgZ*, ΔPA14_00130, ΔPA14_60970, ΔPA14_27930, ΔPA14_56180, Δ*alg44*, ΔPA14_25420 and Δ*pilZ* in swim agar without (grey) and with (black) 1% ethanol after 16 h. Black arrows indicate the candidate mutants of interest (shaded) that were least responsive to ethanol. (B) Swim zone diameter of *P. aeruginosa* PA14 wild type, Δ*pilZ*, Δ*flgZ* and Δ*pilZ*Δ*flgZ* in swim agar without (grey) and with (black) 1% ethanol after 16 h. Error bars indicate standard deviations, n=4 replicates. Common letters indicate samples that are not significantly different, different letters indicate significant differences (*P*-value ˂0.05) as determined by Two-way ANOVA with multiple comparisons.

### Ethanol-dependent motility repression is mediated via the MotAB stator set

Since the PilZ-domain proteins PilZ and FlgZ have been linked to c-di-GMP-dependent decreases in motility mediated by flagellar stators in *P. aeruginosa* and other species (22, 24, 28, 51, 52), we postulated that flagellar stators may also be involved in the response to ethanol. *P. aeruginosa* has two stator sets, MotAB and MotCD (38, 53). Specifically, FlgZ, upon c-di-GMP binding, has been implicated in the sequestration of flagellar motor protein MotC and mislocalization of MotD, which results in loss of motility due to increased incorporation of MotAB which cannot support swimming in many environments such as on swarm agar (22, 24).

In line with the hypothesis that PilZ-domain proteins interact with flagellar stators to reduce motility in the presence of ethanol, the Δ*motAB* mutant had no observable change in motility in the presence of ethanol versus control cultures (Fig. 4). Also consistent with previous reports, the Δ*motCD* mutant displayed a swim zone diameter that was ~90% less than that of wild-type cells grown in the absence of ethanol (24) (Fig. 4). Overall, these data support the conclusion that the MotAB stator set is required for ethanol-dependent swim repression.

**FIG. 4.**
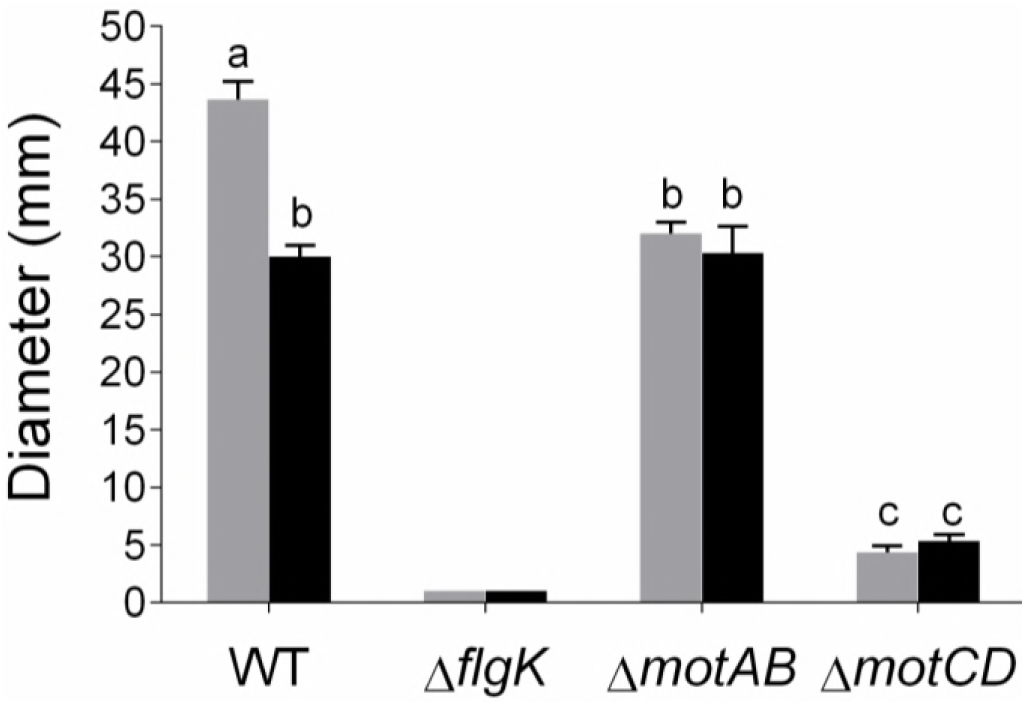
Ethanol effects on swimming motility require the MotAB flagellar stator set. Swim zone diameter of *P. aeruginosa* PA14 wild type, Δ*flgK*, Δ*motAB* and Δ*motCD* in swim agar without (grey) and with (black) 1% ethanol after 16 h. Error bars indicate standard deviations, n=4 replicates. Same letters indicate samples that are not significantly different and different letters have significant differences (*P*-value <0.05) as determined by Multiple t-test corrected using the Holm-Sidak method.

### Ethanol-mediated motility repression requires PilY1 and the PilMNOP Type 4 pili alignment complex

PilY1 has been shown to be a surface-sensing protein required for decreased motility and stimulation of biofilm pathways in cells upon contact with a surface (23, 54) (Fig 5A). PilY1, in conjunction with the type 4 pili (T4P) alignment complex, PilMNOP (54), functions upstream of SadC (23), FlgZ (22) and the MotAB stator (23) to regulate swarming motility in *P. aeruginosa* by controlling the production of and the response to c-di-GMP. Thus, we tested the roles of PilY1 and PilMNOP in ethanol-dependent swimming motility repression. In contrast to the wild-type strain, the Δ*pilY1* mutant did not show decreased motility when ethanol was added to the medium (Fig. 5B). Instead, the Δ*pilY1* mutant showed a reproducible and significant increase in the swim zone diameter when ethanol was added to the medium (Fig. 5B), and a wild-type copy of the *pilY1* gene complemented this phenotype (Fig. 5C). Moreover, a mutant lacking *pilMNOP* showed no ethanol-dependent reduction in swimming motility (Fig. 5B). Though PilY1 and PilMNOP were required for the motility decrease in the presence of ethanol, they were not required for the stimulation of global c-di-GMP levels in planktonic cells (Fig. 5D). These data indicate that PilY1 and PilMNOP are involved in the decreased motility caused by ethanol, and suggest that this may be independent of changing global pools of c-di-GMP.

**FIG. 5.**
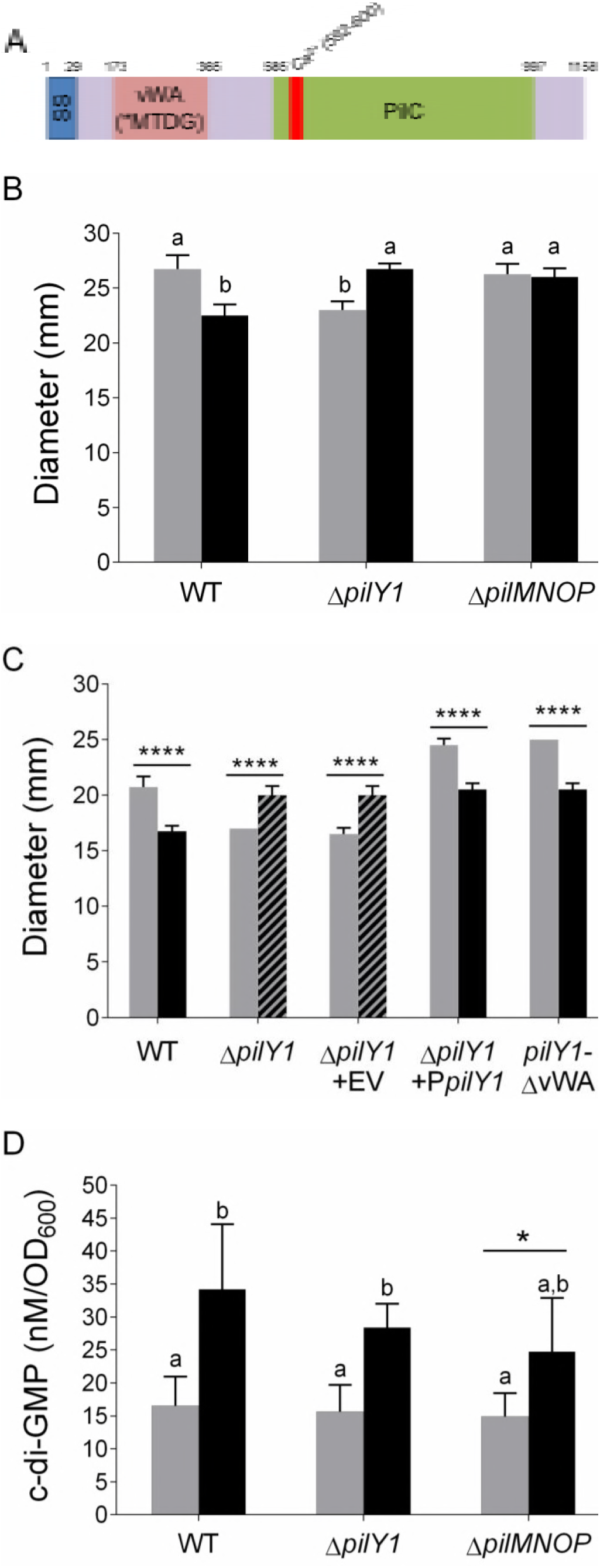
PilY1 and PilMNOP are necessary for ethanol effects on swim zone diameter but not ethanol-dependent c-di-GMP increase. (A) Schematic of the PilY1 protein showing the amino acid positions of the signal sequence (SS), von Willerbrand A factor domain (vWA), calcium binding domain (red), and the PilC domain (green). (B) Swim zone diameter of *P. aeruginosa* PA14 wild type, Δ*pilY1* and Δ*pilMNOP* in swim agar without (grey) and with (black) 1% ethanol measured after 16 h. Error bars indicate standard deviations, n=4 replicates. (C) Swim zone diameter of *P. aeruginosa* PA14 wild type, Δ*pilY1*, Δ*pilY1* with an empty vector (EV), with a plasmid-borne *pilY1* (P*pilY1*), or with a plasmid-borne *pilY1* with the vWA domain deleted (*pilY1*-ΔvWA) in an arabinose-inducible expression plasmid in swim agar without (grey) and with (black) 1% ethanol measured after growth for 16 h. 0.05% arabinose was added to the medium. Error bars indicate standard deviations, n=4 replicates. Hashed bars indicate mutants that swim more in ethanol than in control cultures. (D) Quantification of c-di-GMP levels in *P. aeruginosa* PA14 wild type, Δ*pilY1* and Δ*pilMNOP* grown in liquid M63 medium without (grey) and with (black) 1% ethanol for 16 h. Error bars indicate standard deviations, n=8 replicates. *, *P*-value <0.05; ****, *P*-value <0.0001 as determined by Two-way ANOVA with multiple comparisons. Same letters indicate no significant differences, different letters indicate samples that are significantly different (*P*-value <0.05).

### Type 4 pili are not required for the reduction in swimming motility in response to ethanol

PilY1 is necessary for T4P activity (55), and thus we sought to determine if PilY1- and PilMNOP-dependent reduction in flagellar motility by ethanol was due to a decrease in T4P activity. Two pieces of evidence argue against a role for the T4P in ethanol-mediated effects on motility. First, a Δ*pilA* mutant (which lacks pili) still showed the same level of motility and responsiveness to ethanol (Fig. S5A) when compared to wild-type cells. Secondly, ethanol did not reduce twitching motility in wild-type cells (Fig. S5B); rather, a small but significant increase in twitch zone diameter was observed in cultures with ethanol. These data suggest that the ethanol effects on swimming motility are not due to changes in T4P function.

### Previously described elements involved in PilY1 activation were dispensable for the ethanol-dependent reduction in motility

We sought to determine if previously described factors involved in PilY1 activation were involved in the ethanol response. Previous studies had shown that *pilY1* transcription is regulated by PilJ, a component of the Pil-Chp pathway, in response to surface engagement (54) through stimulation of cAMP production (56) upon surface contact (54). The *cyaAB* genes, which encode adenylate cyclases responsible for cAMP production by *P. aeruginosa*, were also implicated in PilY1 activation (54). We found that the Δ*pilJ* and Δ*cyaAB* mutants, though hyper motile in control cultures, both exhibited motility repression in response to ethanol (p<0.0001, Fig. S6). These data indicated that the upstream cAMP signal previously shown to be required for *pilY1* transcription, upon surface contact, was not required for the PilY1-dependent changes in motility in response to ethanol.

The von Willerbrand factor A (vWA) domain of PilY1, depicted in the schematic in Fig. 5A, is necessary for surface-associated swarming motility repression (23). To probe whether the same domain of PilY1 required for surface-sensing was also necessary for ethanol responsiveness, we used a strain where a mutated *pilY1* with the vWA domain deleted (*pilY1*-ΔvWA) was placed at the native *pilY1* locus. The *pilY1-*ΔvWA strain was still responsive to ethanol-dependent motility repression (25 mm and 20.5 ± 0.3 mm in the absence and presence of ethanol, respectively (p<0.0001; Fig. 5C). These data indicate that the vWA domain of PilY1 is dispensable for ethanol-mediated swim repression.

### Increased flagellar reversal frequency in the presence of ethanol does not cause ethanol-dependent motility repression

*P. aeruginosa* is a monotrichous flagellated bacterium whose flagellar motility is governed by a run-reverse pattern (32, 57) rather than the run-tumble pattern in organisms with peritrichous flagella like *E. coli* (58). *P. aeruginosa* directional movement is due to a change in the flagellar rotation (clockwise or counterclockwise) and this rotation change is called a ‘reversal’. The frequency of reversals can impact the area covered since *P. aeruginosa* must slow its normal speed from 40–55 µm/sec (38, 53, 57) to as low as 15 µm/sec immediately before a reversal (57). To determine if the decrease in swim zone diameter in ethanol was a result of a change in reversal frequencies, this parameter was measured in the absence and presence of 1% ethanol.

Cells from mid-exponential phase cultures were treated with ethanol for 15 min prior to measurement of the reversal frequency. *P. aeruginosa* strain PA14 wild type showed a 4.8-fold increase in its reversal frequency in the presence of ethanol, going from 4.6 ± 4.1 to 22 ± 9.1 reversals/ 10 s (p<0.0001; Fig. S7A). Similarly, the Δ*sadC*Δ*gcbA*, Δ*pilY1*, and Δ*pilMNOP* mutants also showed significant 2.8-fold, 3.4-fold, and 2.7-fold (p<0.0001) increases in reversal frequencies upon the inclusion of ethanol in the medium (Fig. S7A-B). There were no significant differences between the wild type and the mutants in either the control (except for Δ*sadC*Δ*gcbA*) or ethanol conditions (Fig. S7A-B). These data indicated that in the presence of ethanol, *P. aeruginosa* had a higher rate of flagellar reversals than in control conditions, but this change did not account for the SadC/GcbA-, PilY1-, or PilMNOP-dependent suppression of motility in the presence of ethanol.

### Ethanol rapidly increases the sub-population of immobile cells in swim agar, in a PilY1-, PilMNOP-, FlgZ-, PilZ-, and MotAB-dependent manner

We next observed the behavior of single cells in swim agar in the absence and presence of 1% ethanol in order to better understand how ethanol affected the macroscopic swim zone size. To do this experiment, we exposed exponentially growing cells to swim agar without and with ethanol for 30 min followed by the acquisition of 8 s time-lapse movies to visualize cellular behavior as outlined in Fig. 6A.

**FIG. 6.**
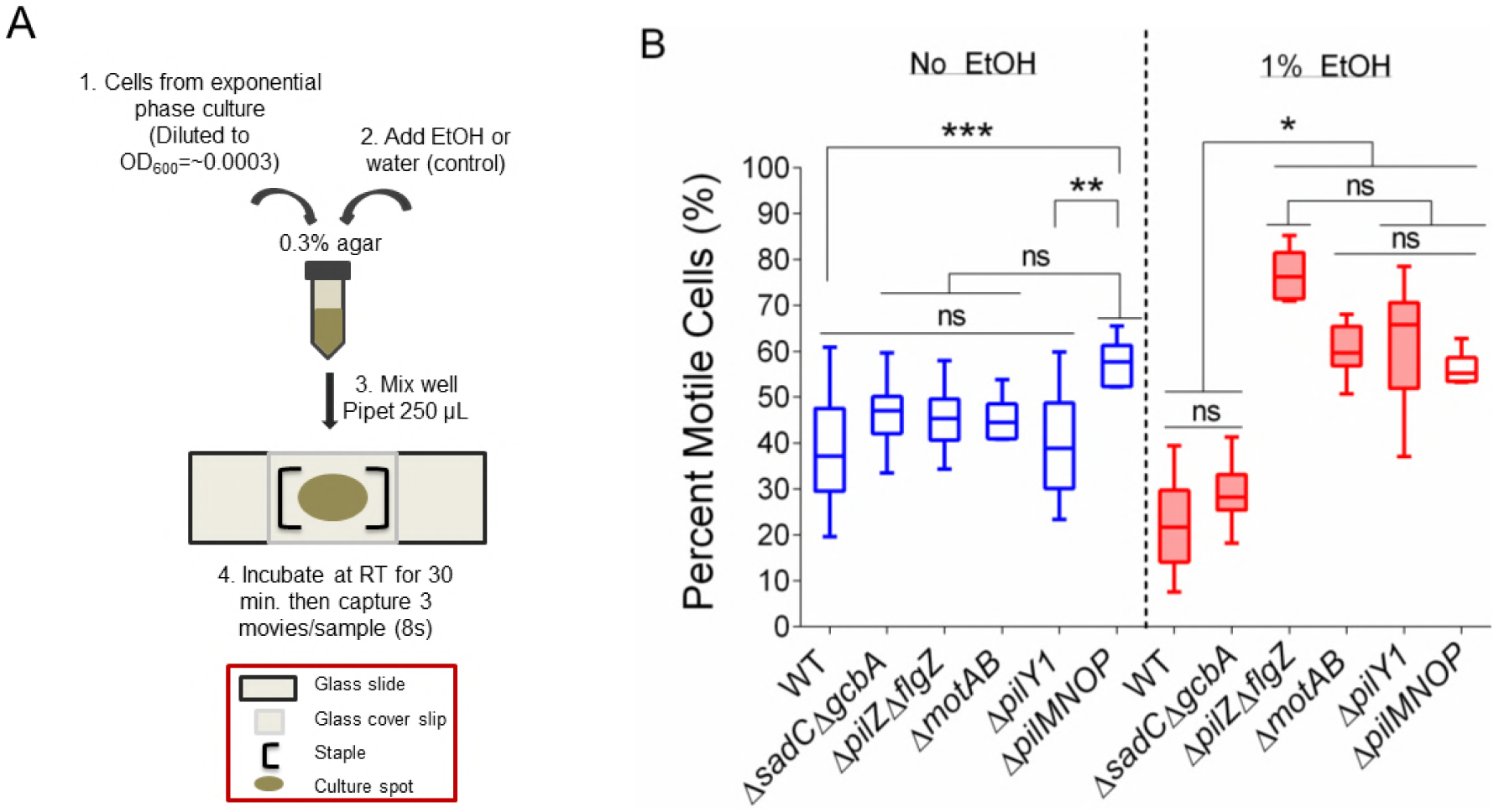
Ethanol decreases the number of motile cells in swim agar, in a manner dependent on PilY1, PilMNOP, FlgZ, PilZ, and MotAB. (A) Schematic of agar motility assay treated with water (control) or 1% ethanol in swim agar. Sample was mixed well then 250 µL was pipetted onto a glass slide and staples were used to create a chamber using a glass coverslip. Sample was then incubated at room temperature for 30 min, and then imaged as outlined in the methods. (B) Agar motility assay of *P. aeruginosa* PA14 wild type, Δ*sadC*Δ*gcbA*, Δ*pilZ*Δ*flgZ*, Δ*motAB*, Δ*pilY1* and Δ*pilMNOP* in swim agar without (blue) and with (red) 1% ethanol after 30 min. Three time lapse (8 s) movies of the cells in the agar matrix, for each sample, were captured. Plotted is the average population percentage of the motile subpopulation in each movie. Error bars represent the maximum and minimum data point, n≥6 replicate movies. Shaded boxes represent the ethanol samples that are significantly different from their controls. *, *P*-value <0.05; **, *P*-value <0.01; ***, *P*-value <0.001; ns, not significant as determined by One-Way ANOVA with multiple comparisons.

We first noted that when the fraction of motile cells in the control and ethanol-treated samples were compared for each mutant, all except Δ*pilMNOP* were statistically different (Fig. 6B). We also noted that in the control cultures, Δ*pilMNOP* was significantly higher than wild type and Δ*pilY1* in the same condition (Fig. 6B). For *P. aeruginosa* PA14 wild type cells swimming in swim agar, we observed a decrease in the fraction of motile cells in the presence of ethanol within the 8 s time interval analyzed (38 ± 10.5% motile (control) and 22 ± 8.8% motile (ethanol); p≤0.05; Fig. 6B). In contrast, most of the mutants that did not show a smaller swim zone in response to ethanol also did not show a reduction in the fraction of motile cells when ethanol was in the medium. The Δ*pilMNOP* mutant, for example, had a similar proportion of motile cells in the absence and presence of ethanol (58 ± 5.1% and 56 ± 3.6%, respectively; Fig. 6B). Interestingly, the Δ*pilY1*, Δ*pilZ*Δ*flgZ*, and Δ*motAB* cells showed an increase in the fraction of motile cells in the presence of ethanol (Fig. 6B) which also mirrored the observation that the Δ*pilY1* strain had a larger swim zone size in the presence of ethanol (Fig. 5B). Of the mutants that were resistant to the effects of ethanol in the macroscopic swim zone assay, only the Δ*sadC*Δ*gcbA* double mutant was not significantly different from the wild-type strain (46 ± 6.4% motile (control) and 29 ± 6.4% motile (ethanol); Fig. 6B). These data suggest that in response to ethanol, *P. aeruginosa* exhibits an increase in the periods of immobility or decrease in the fraction of cells swimming in the swim agar, and that this response is dependent on PilY1, PilMNOP, FlgZ, PilZ, and MotAB proteins. The observation that the Δ*sadC*Δ*gcbA* double mutant behaves like the wild type in the microscopic assay may suggest functional redundancy with other c-di-GMP enzymes or that c-di-GMP levels affect motility by a mechanism distinct from that of the PilY1-PilMNOP-FlgZ/PilZ-MotAB pathway.

### Ethanol inhibits swarming motility

In addition to its effects on planktonic cells and motility in swim agar, ethanol also inhibits flagellum-dependent swarming motility on agar surfaces (Fig. 7A) (5). We found that PilY1, the PilMNOP alignment complex, the PilZ and FlgZ proteins, and the MotAB stators, which were all required to increase the fraction of sessile cells in medium with ethanol, were also necessary for full suppression of flagellar-mediated swarming motility on the surface of 0.5% agar in the presence of ethanol (Fig. 7A-D). Furthermore, while the vWA domain of PilY1 has been shown to be important for surface-sensing (23), we found that this domain was not required for ethanol-mediated repression of swarming motility (Fig. 7A).

**FIG. 7.**
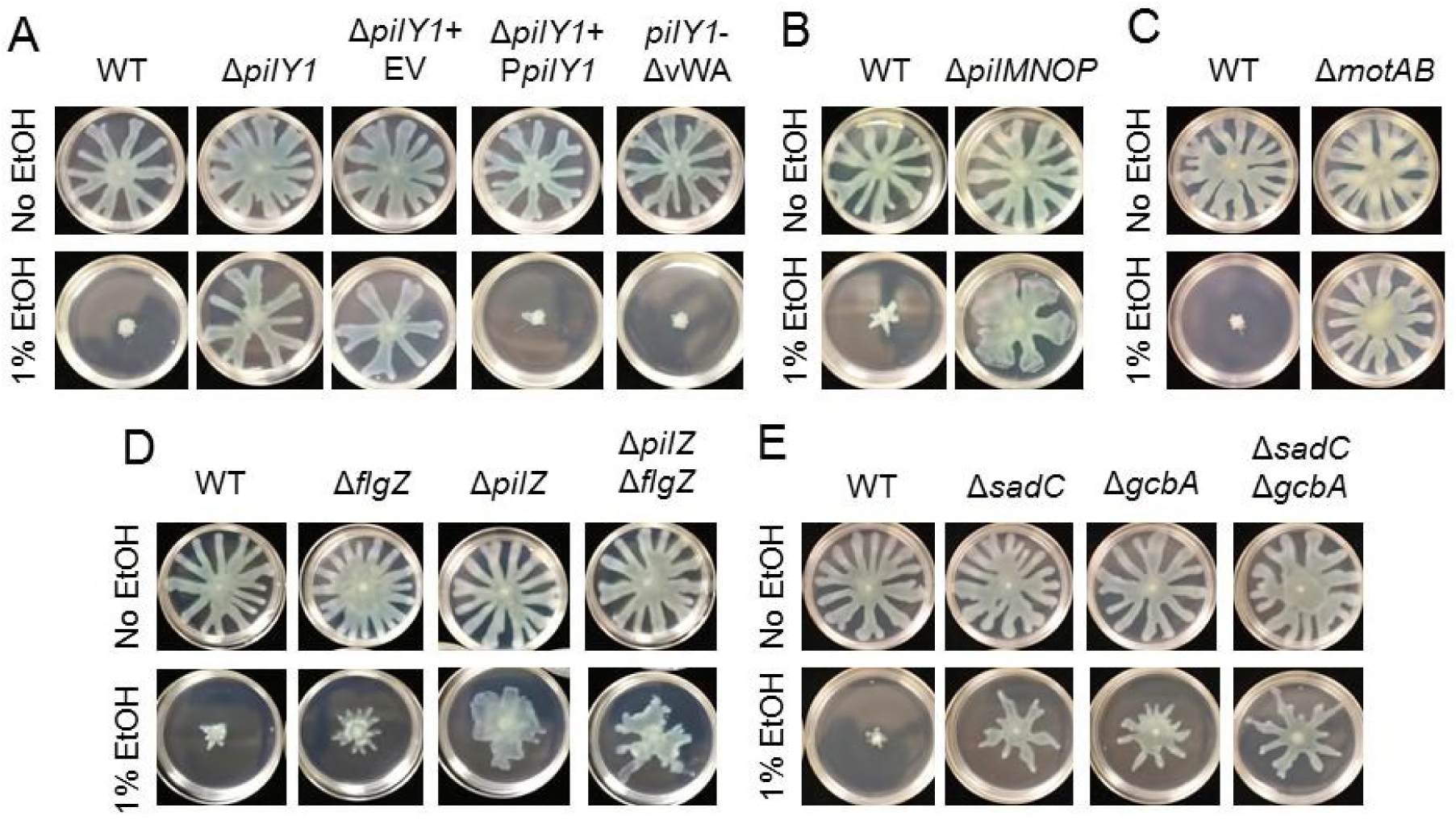
Ethanol effects on swarming motility repression occur using some surface-sensing components as well as components independent of surface sensing. Representative images of swarming motility assays of (A) *P. aeruginosa* PA14 wild type, Δ*pilY1*, Δ*pilY1* with an empty vector (EV) or with a plasmid that enables arabinose-inducible expression *pilY1* (P*pilY1*) or *pilY1* without a vWA domain (*pilY1*-ΔvWA), (B) *P. aeruginosa* PA14 wild type and Δ*pilMNOP*, (C) *P. aeruginosa* PA14 wild type and Δ*motAB*, (D) *P. aeruginosa* PA14 wild type, Δ*flgZ*, Δ*pilZ*, and Δ*pilZ*Δ*flgZ*, and (E) *P. aeruginosa* PA14 wild type, Δ*sadC*, Δ*gcbA*, and Δ*sadC*Δ*gcbA* in M63 medium with 0.5% agar (swarm agar) without and with 1% ethanol and grown for 16 h. Images are representative of observed phenotypes, n=4 per experiment and each experiment was performed 3–5 times.

Consistent with our observation that the Δ*sadC*, Δ*gcbA*, and Δ*sadC*Δ*gcbA* mutants were resistant to the effects of ethanol on motility in the swim agar assay, these mutants were also less responsive to ethanol in the surface-associated swarming motility assay (Fig. 7E). Together, these data suggest that common factors are involved in the repression of flagellar motility in planktonic cells as well as in cells on a surface.

## Discussion

Here we present a model (Fig. 8) in which ethanol leads to decreased flagellar motility in *P. aeruginosa*. The down-regulation of flagellar motility by ethanol is in line with our previous work and the work of others that together shows that ethanol (1%) increases biofilm formation on abiotic and biotic surfaces (5, 42). We identified a pathway that involves PilY1, the T4P alignment complex (PilMNOP), two PilZ-domain proteins (FlgZ and PilZ) and the stator MotAB, all of which are required for ethanol-mediated down-regulation of swimming motility in both macroscopic and microscopic swim motility assays and for flagellar-mediated swarming motility on a surface. Microscopic observations of cells in swim agar, a medium that is widely used to assess chemotaxis and swimming motility (18, 59–61), showed that ethanol decreased the fraction of cells that were motile during the image capture period and that this change in behavior required all of the components of the PilY1 pathway outlined above. This same PilY1/PilMNOP/PilZ/FlgZ/MotAB pathway was shown previously to play a key role in surface sensing and early biofilm formation by *P. aeruginosa* (22–24, 54). FlgZ, a homolog of *E. coli* YcgR, has been shown to regulate flagellar motility by directly interacting with the flagellar motor proteins, thereby behaving like a ‘brake’ for flagellar rotation (22, 51, 62). The finding that ethanol reduces flagellar motility in cells in the soft agar suspension may provide more insight into the mechanism by which external or membrane-localized signals transduced by PilY1 and PilMNOP cause FlgZ, and perhaps PilZ, to act to affect flagellar rotation.

**FIG. 8.**
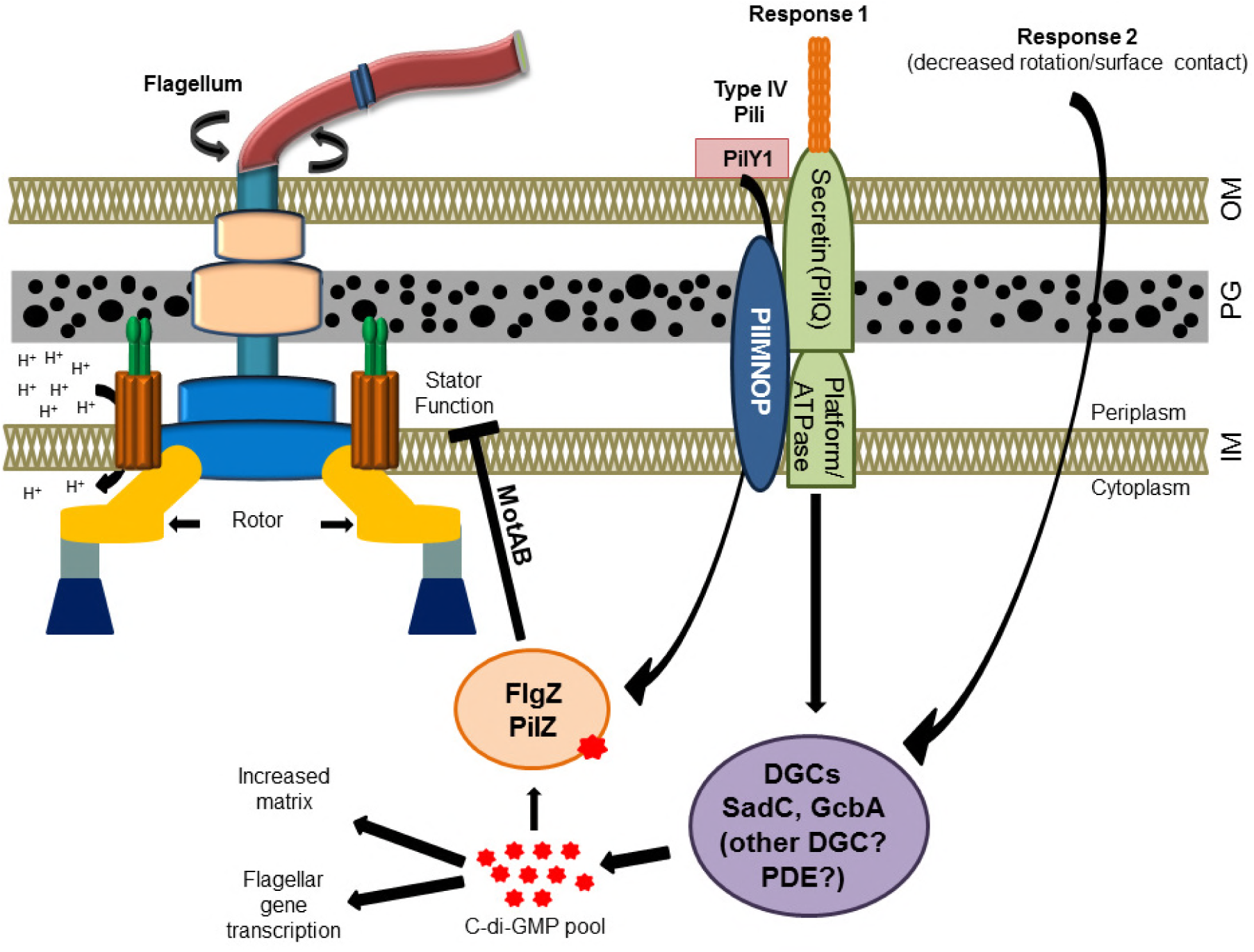
Model for the effects of ethanol on *Pseudomonas aeruginosa* motility. When free swimming *P. aeruginosa* encounters ethanol in a liquid environment, it quickly responds by changing its motility. One response (1) occurs via PilY1 and the alignment complex (PilMNOP). This motility repression requires two PilZ domain proteins, PilZ and FlgZ, as well as flagellar stator set, MotAB. We propose that this signal causes the flagellar machinery to ‘brake’, resulting in a decrease in the number of cells that are motile. The second response (2) involves a SadC and GcbA c-di-GMP production-dependent change that may also involve other c-di-GMP metabolic proteins. C-di-GMP may further activate the PilZ domain proteins. Together, these responses repress flagellar motility in swim agar conditions and a soft agar that supports swarming motility.

We also observed an ethanol-dependent increase in global pools of c-di-GMP in planktonic cells that required the activity of the diguanylate cyclases (DGCs) SadC and GcbA. In the plate-based swim assay, the ∆*sadC*∆*gcbA* double mutant no longer showed an ethanol-mediated reduction in swimming motility. However, this double mutant behaved similarly to the wild type in the microscopic swim assay; we do not fully understand the basis for this discrepancy. The differences in swim zone diameters may be the consequence of a combination of factors that influence flagellar motility in different ways over the course of hours, while the short-term microscopic assay may only assess a subset of the early effects of ethanol. Previous studies have highlighted the distinct roles that SadC and GcbA play during the different stages of biofilm formation. GcbA, for example, was implicated in c-di-GMP production only in planktonic cells or cells initiating biofilm formation or dispersing from a mature biofilm (21, 25, 63). SadC, on the other hand, is implicated in biofilm initiation and maturation (7, 21). It is also important to note that the ∆*sadC*∆*gcbA* double mutant still showed a significant, albeit reduced, increase in c-di-GMP in the presence of ethanol (Fig. 2B), and thus other c-di-GMP metabolizing enzymes may be involved in ethanol-mediated swimming repression, a finding consistent with our initial genetic screen. In addition, c-di-GMP metabolizing enzymes can affect target protein activities either by not altering global pools of c-di-GMP (local signaling) or by increasing global levels of c-di-GMP by overexpression of a DGC (6, 7, 23, 25) or disruption of a PDE (6, 22, 23). It is also possible that the status of global c-di-GMP is important and may alter how *P. aeruginosa* responds to ethanol. Future studies will dissect the contributions of other c-di-GMP metabolic activities on ethanol-induced effects on motility and other c-di-GMP controlled processes.

A key question is how does *P. aeruginosa* PilY1 and PilMNOP, which are localized to the membrane and extracytoplasmic space, contribute to the repression of motility in the presence of ethanol? We showed that the N-terminal vWA domain of PilY1 is dispensable for responding to ethanol, implicating the C-terminal domain of this protein as key for the observed ethanol response. The C-terminus of the PilY1 protein has a seven-bladed, modified β-propeller structure that shares structural similarity to the quinohemoprotein alcohol dehydrogenase from *Comamonas testosteroni* (55). An attractive hypothesis is that PilY1 has the ability to bind ethanol or the co-factor required for its catabolism. Alternatively, alcohols such as ethanol have been implicated in membrane perturbation (64–66). For example, in *E. coli*, proteomics analysis during ethanol stress, using 4% (684 mM) ethanol, revealed an induction of the general stress response, a 15% increase in membrane fluidity, and the induction of mechanosensitive channels that are active during osmotic stress (66). In another publication, Cao *et. al*. also showed that in *E. coli*, the effect of 2.5–5% (428–855 mM) ethanol resulted in increased ROS stress, reduced peptidoglycan, and a decrease in the proton gradient that might be explained by increased membrane fluidity (65). Cao *et. al*. also showed that evolution of *E. coli* on ethanol resulted in the correction of all the transient changes listed above, due to increased mutation rates (65). It is possible that this membrane perturbation also occurs in *P. aeruginosa* upon ethanol exposure and might induce structural changes to proteins associated with the cell membrane (PilY1, PilMNOP and SadC) that would then play a role in their activation. It is also possible that there are unknown regulators that are activated by ethanol and induce the activity of the pathway described above.

We also found that *P. aeruginosa* flagellar reversal frequency was significantly increased in the presence of ethanol, but this response did not depend on any of the proteins that we tested. Previous studies have indicated that an increase in reversal frequency increases the cell’s ability to move more efficiently through soft agar (7, 67). Additionally, *P. aeruginosa* and related bacteria that utilize a run-reverse-turn trajectory, spend equal time going clockwise or counterclockwise with variation in their pause duration in order to turn at different angles to maximize space exploration (32). Since chemotaxis involves the modulation of reversal frequencies (57), these data may suggest that ethanol also affect chemotaxis in *P. aeruginosa*. More work is required to determine if ethanol influences positive or negative chemotactic pathways. Together, our data suggest that, while ethanol reduces the fraction of motile *P. aeruginosa* cells within a given time interval, these motile cells can navigate a viscous environment more efficiently in order to remain in the local space of the ethanol-producing microbes.

To conclude, our findings indicate that ethanol triggers a complex response that modulates behaviors related to biofilm initiation in order to facilitate the transition from being motile to being sessile. Therefore, the effects of ethanol on microbes at concentrations much lower than those used for the purpose of sterilization is of interest in the context of biofuel production, microbial remediation of industrial waste, and the activity of naturally occurring communities in the environment and those in association with humans. Future studies will determine if ethanol’s effects on *P. aeruginosa* motility contributes to the stimulation of biofilm formation and if the effects of ethanol on motility and biofilm formation in other Gram-negative species, like *Acinetobacter baumannii* (4), occurs through a common pathway.

## Acknowledgements

Research reported in this publication was also supported by grants from the National Institutes of Health to D.A.H. (R01 GM108492 to D.A.H) and to G.A.O. (R37 AI83256), a pilot project from STANTO19R0, and NSF 1458359 (D.A.H. and K.A.L.). Support for C.E.H. came in part from 5T32AI007519. Additional support was provided by the NCI Cancer Center Support Grant, 5P30 CA023108, through the Molecular Biology Shared Resource, and NIGMS P20GM113132 through the Molecular Interactions and Imaging Core (MIIC). We also thank Dr. Karin Sauer for sharing reagents, Emily L. Dolben for assistance with early experiments, Alan J. Collins for designing the chamber slide setup used in the macroscopic agar motility assays, and Dr. Dae Gon Ha for making the *gcbA* complementation construct.

## References

1. Ranger, C. M., Biedermann, P. H. W., Phuntumart, V., Beligala, G. U., Ghosh, S., Palmquist, D. E., Mueller, R., Barnett, J., Schultz, P. B., Reding, M. E. and Benz, J. P. 2018. Symbiont selection via alcohol benefits fungus farming by ambrosia beetles. Proc Natl Acad Sci U S A 115:4447–4452.

2. Lovett, J. L., Marchesini, N., Moreno, S. N. and Sibley, L. D. 2002. *Toxoplasma gondii* microneme secretion involves intracellular Ca(2+) release from inositol 1,4,5-triphosphate (IP(3))/ryanodine-sensitive stores. J Biol Chem 277:25870–6.

3. Carruthers, V. B., Moreno, S. N. and Sibley, L. D. 1999. Ethanol and acetaldehyde elevate intracellular [Ca2+] and stimulate microneme discharge in *Toxoplasma gondii*. Biochem J 342 (Pt 2):379–86.

4. Nwugo, C. C., Arivett, B. A., Zimbler, D. L., Gaddy, J. A., Richards, A. M. and Actis, L. A. 2012. Effect of ethanol on differential protein production and expression of potential virulence functions in the opportunistic pathogen *Acinetobacter baumannii*. PLoS One 7:e51936.

5. Chen, A. I., Dolben, E. F., Okegbe, C., Harty, C. E., Golub, Y., Thao, S., Ha, D. G., Willger, S. D., O’Toole, G. A., Harwood, C. S., Dietrich, L. E. and Hogan, D. A. 2014. *Candida albicans* ethanol stimulates *Pseudomonas aeruginosa* WspR-controlled biofilm formation as part of a cyclic relationship involving phenazines. PLoS Pathog 10:e1004480.

6. Merighi, M., Lee, V. T., Hyodo, M., Hayakawa, Y. and Lory, S. 2007. The second messenger bis-(3’-5’)-cyclic-GMP and its PilZ domain-containing receptor Alg44 are required for alginate biosynthesis in *Pseudomonas aeruginosa*. Mol Microbiol 65:876–95.

7. Merritt, J. H., Brothers, K. M., Kuchma, S. L. and O’Toole, G. A. 2007. SadC reciprocally influences biofilm formation and swarming motility via modulation of exopolysaccharide production and flagellar function. J Bacteriol 189:8154–64.

8. Montuschi, P., Paris, D., Melck, D., Lucidi, V., Ciabattoni, G., Raia, V., Calabrese, C., Bush, A., Barnes, P. J. and Motta, A. 2012. NMR spectroscopy metabolomic profiling of exhaled breath condensate in patients with stable and unstable cystic fibrosis. Thorax 67:222–8.

9. Montuschi, P., Paris, D., Montella, S., Melck, D., Mirra, V., Santini, G., Mores, N., Montemitro, E., Majo, F., Lucidi, V., Bush, A., Motta, A. and Santamaria, F. 2014. Nuclear magnetic resonance-based metabolomics discriminates primary ciliary dyskinesia from cystic fibrosis. Am J Respir Crit Care Med 190:229–33.

10. Worlitzsch, D., Tarran, R., Ulrich, M., Schwab, U., Cekici, A., Meyer, K. C., Birrer, P., Bellon, G., Berger, J., Weiss, T., Botzenhart, K., Yankaskas, J. R., Randell, S., Boucher, R. C. and Doring, G. 2002. Effects of reduced mucus oxygen concentration in airway *Pseudomonas* infections of cystic fibrosis patients. J Clin Invest 109:317–25.

11. Bos, L. D., Meinardi, S., Blake, D. and Whiteson, K. 2016. Bacteria in the airways of patients with cystic fibrosis are genetically capable of producing VOCs in breath. J Breath Res 10:047103.

12. Wolak, J. E., Esther, C. R., Jr. and O’Connell, T. M. 2009. Metabolomic analysis of bronchoalveolar lavage fluid from cystic fibrosis patients. Biomarkers 14:55–60.

13. Sofia, M., Maniscalco, M., de Laurentiis, G., Paris, D., Melck, D. and Motta, A. 2011. Exploring airway diseases by NMR-based metabonomics: a review of application to exhaled breath condensate. J Biomed Biotechnol 2011:403260.

14. Merritt, J. H., Ha, D. G., Cowles, K. N., Lu, W., Morales, D. K., Rabinowitz, J., Gitai, Z. and O’Toole, G. A. 2010. Specific control of *Pseudomonas aeruginosa* surface-associated behaviors by two c-di-GMP diguanylate cyclases. MBio 1.

15. Stelitano, V., Giardina, G., Paiardini, A., Castiglione, N., Cutruzzola, F. and Rinaldo, S. 2013. C-di-GMP hydrolysis by Pseudomonas aeruginosa HD-GYP phosphodiesterases: analysis of the reaction mechanism and novel roles for pGpG. PLoS One 8:e74920.

16. Kuchma, S. L., Brothers, K. M., Merritt, J. H., Liberati, N. T., Ausubel, F. M. and O’Toole, G. A. 2007. BifA, a cyclic-Di-GMP phosphodiesterase, inversely regulates biofilm formation and swarming motility by Pseudomonas aeruginosa PA14. J Bacteriol 189:8165–78.

17. Kazmierczak, B. I., Lebron, M. B. and Murray, T. S. 2006. Analysis of FimX, a phosphodiesterase that governs twitching motility in *Pseudomonas aeruginosa*. Mol Microbiol 60:1026–43.

18. Ha, D. G., Richman, M. E. and O’Toole, G. A. 2014. Deletion mutant library for investigation of functional outputs of cyclic diguanylate metabolism in *Pseudomonas aeruginosa* PA14. Appl Environ Microbiol 80:3384–93.

19. Giacalone, D., Smith, T. J., Collins, A. J., Sondermann, H., Koziol, L. J. and O’Toole, G. A. 2018. Ligand-mediated biofilm formation via enhanced physical interaction between a diguanylate cyclase and its receptor. MBio 9.

20. Pei, J. and Grishin, N. V. 2001. GGDEF domain is homologous to adenylyl cyclase. Proteins 42:210–6.

21. Valentini, M. and Filloux, A. 2016. Biofilms and cyclic di-GMP (c-di-GMP) signaling: lessons from *Pseudomonas aeruginosa* and other bacteria. J Biol Chem 291:12547–55.

22. Baker, A. E., Diepold, A., Kuchma, S. L., Scott, J. E., Ha, D. G., Orazi, G., Armitage, J. P. and O’Toole, G. A. 2016. PilZ domain protein FlgZ mediates cyclic di-GMP-dependent swarming motility control in *Pseudomonas aeruginosa*. J Bacteriol 198:1837–46.

23. Kuchma, S. L., Ballok, A. E., Merritt, J. H., Hammond, J. H., Lu, W., Rabinowitz, J. D. and O’Toole, G. A. 2010. Cyclic-di-GMP-mediated repression of swarming motility by *Pseudomonas aeruginosa*: the *pilY1* gene and its impact on surface-associated behaviors. J Bacteriol 192:2950–64.

24. Kuchma, S. L., Delalez, N. J., Filkins, L. M., Snavely, E. A., Armitage, J. P. and O’Toole, G. A. 2015. Cyclic di-GMP-mediated repression of swarming motility by *Pseudomonas aeruginosa* PA14 requires the MotAB stator. J Bacteriol 197:420–30.

25. Petrova, O. E., Cherny, K. E. and Sauer, K. 2014. The *Pseudomonas aeruginosa* diguanylate cyclase GcbA, a homolog of P. fluorescens GcbA, promotes initial attachment to surfaces, but not biofilm formation, via regulation of motility. J Bacteriol 196:2827–41.

26. Wolfe, A. J. and Visick, K. L. 2008. Get the message out: cyclic-Di-GMP regulates multiple levels of flagellum-based motility. J Bacteriol 190:463–75.

27. Waters, C. M. 2013. Bacterial wheel locks: extracellular polysaccharide inhibits flagellar rotation. J Bacteriol 195:409–10.

28. Zorraquino, V., Garcia, B., Latasa, C., Echeverz, M., Toledo-Arana, A., Valle, J., Lasa, I. and Solano, C. 2013. Coordinated cyclic-di-GMP repression of *Salmonella* motility through YcgR and cellulose. J Bacteriol 195:417–28.

29. Choy, W. K., Zhou, L., Syn, C. K., Zhang, L. H. and Swarup, S. 2004. MorA defines a new class of regulators affecting flagellar development and biofilm formation in diverse *Pseudomonas* species. J Bacteriol 186:7221–8.

30. Ko, M. and Park, C. 2000. Two novel flagellar components and H-NS are involved in the motor function of *Escherichia coli*. J Mol Biol 303:371–82.

31. Blair, K. M., Turner, L., Winkelman, J. T., Berg, H. C. and Kearns, D. B. 2008. A molecular clutch disables flagella in the *Bacillus subtilis* biofilm. Science 320:1636–8.

32. Qian, C., Wong, C. C., Swarup, S. and Chiam, K. H. 2013. Bacterial tethering analysis reveals a “run-reverse-turn” mechanism for *Pseudomonas* species motility. Appl Environ Microbiol 79:4734–43.

33. Paul, K., Nieto, V., Carlquist, W. C., Blair, D. F. and Harshey, R. M. 2010. The c-di-GMP binding protein YcgR controls flagellar motor direction and speed to affect chemotaxis by a “backstop brake” mechanism. Mol Cell 38:128–39.

34. Kojima, S. and Blair, D. F. 2001. Conformational change in the stator of the bacterial flagellar motor. Biochemistry 40:13041–50.

35. Braun, T. F. and Blair, D. F. 2001. Targeted disulfide cross-linking of the MotB protein of *Escherichia coli*: evidence for two H(+) channels in the stator complex. Biochemistry 40:13051–9.

36. Blair, D. F. 2003. Flagellar movement driven by proton translocation. FEBS Lett 545:86–95.

37. Toutain, C. M., Caizza, N. C., Zegans, M. E. and O’Toole, G. A. 2007. Roles for flagellar stators in biofilm formation by *Pseudomonas aeruginosa*. Res Microbiol 158:471–7.

38. Doyle, T. B., Hawkins, A. C. and McCarter, L. L. 2004. The complex flagellar torque generator of *Pseudomonas aeruginosa*. J Bacteriol 186:6341–50.

39. Shanks, R. M., Caiazza, N. C., Hinsa, S. M., Toutain, C. M. and O’Toole, G. A. 2006. *Saccharomyces cerevisiae*-based molecular tool kit for manipulation of genes from Gram-negative bacteria. Appl Environ Microbiol 72:5027–36.

40. Caiazza, N. C., Merritt, J. H., Brothers, K. M. and O’Toole, G. A. 2007. Inverse regulation of biofilm formation and swarming motility by *Pseudomonas aeruginosa* PA14. J Bacteriol 189:3603–12.

41. Schindelin, J., Arganda-Carreras, I., Frise, E., Kaynig, V., Longair, M., Pietzsch, T., Preibisch, S., Rueden, C., Saalfeld, S., Schmid, B., Tinevez, J. Y., White, D. J., Hartenstein, V., Eliceiri, K., Tomancak, P. and Cardona, A. 2012. Fiji: an open-source platform for biological-image analysis. Nat Methods 9:676–82.

42. Tashiro, Y., Inagaki, A., Ono, K., Inaba, T., Yawata, Y., Uchiyama, H. and Nomura, N. 2014. Low concentrations of ethanol stimulate biofilm and pellicle formation in *Pseudomonas aeruginosa*. Biosci Biotechnol Biochem 78:178–81.

43. Gorisch, H. 2003. The ethanol oxidation system and its regulation in *Pseudomonas aeruginosa*. Biochim Biophys Acta 1647:98–102.

44. Mern, D. S., Ha, S. W., Khodaverdi, V., Gliese, N. and Gorisch, H. 2010. A complex regulatory network controls aerobic ethanol oxidation in *Pseudomonas aeruginosa*: indication of four levels of sensor kinases and response regulators. Microbiology 156:1505–16.

45. Lee, V. T., Matewish, J. M., Kessler, J. L., Hyodo, M., Hayakawa, Y. and Lory, S. 2007. A cyclic-di-GMP receptor required for bacterial exopolysaccharide production. Mol Microbiol 65:1474–84.

46. Whitney, J. C., Colvin, K. M., Marmont, L. S., Robinson, H., Parsek, M. R. and Howell, P. L. 2012. Structure of the cytoplasmic region of PelD, a degenerate diguanylate cyclase receptor that regulates exopolysaccharide production in *Pseudomonas aeruginosa*. J Biol Chem 287:23582–93.

47. Baraquet, C. and Harwood, C. S. 2016. FleQ DNA binding consensus sequence revealed by studies of FleQ-dependent regulation of biofilm gene expression in *Pseudomonas aeruginosa*. J Bacteriol 198:178–86.

48. Hengge, R. 2009. Principles of c-di-GMP signalling in bacteria. Nat Rev Microbiol 7:263–73.

49. Pratt, J. T., Tamayo, R., Tischler, A. D. and Camilli, A. 2007. PilZ domain proteins bind cyclic diguanylate and regulate diverse processes in *Vibrio cholerae*. J Biol Chem 282:12860–70.

50. Christen, M., Christen, B., Allan, M. G., Folcher, M., Jeno, P., Grzesiek, S. and Jenal, U. 2007. DgrA is a member of a new family of cyclic diguanosine monophosphate receptors and controls flagellar motor function in *Caulobacter crescentus*. Proc Natl Acad Sci U S A 104:4112–7.

51. Ryjenkov, D. A., Simm, R., Romling, U. and Gomelsky, M. 2006. The PilZ domain is a receptor for the second messenger c-di-GMP: the PilZ domain protein YcgR controls motility in enterobacteria. J Biol Chem 281:30310–4.

52. Boehm, A., Kaiser, M., Li, H., Spangler, C., Kasper, C. A., Ackermann, M., Kaever, V., Sourjik, V., Roth, V. and Jenal, U. 2010. Second messenger-mediated adjustment of bacterial swimming velocity. Cell 141:107–16.

53. Toutain, C. M., Zegans, M. E. and O’Toole, G. A. 2005. Evidence for two flagellar stators and their role in the motility of *Pseudomonas aeruginosa*. J Bacteriol 187:771–7.

54. Luo, Y., Zhao, K., Baker, A. E., Kuchma, S. L., Coggan, K. A., Wolfgang, M. C., Wong, G. C. and O’Toole, G. A. 2015. A hierarchical cascade of second messengers regulates *Pseudomonas aeruginosa* surface behaviors. MBio 6.

55. Orans, J., Johnson, M. D., Coggan, K. A., Sperlazza, J. R., Heiniger, R. W., Wolfgang, M. C. and Redinbo, M. R. 2010. Crystal structure analysis reveals *Pseudomonas* PilY1 as an essential calcium-dependent regulator of bacterial surface motility. Proc Natl Acad Sci U S A 107:1065–70.

56. Belete, B., Lu, H. and Wozniak, D. J. 2008. *Pseudomonas aeruginosa* AlgR regulates type IV pilus biosynthesis by activating transcription of the *fimU*-*pilVWXY1Y2E* operon. J Bacteriol 190:2023–30.

57. Cai, Q., Li, Z., Ouyang, Q., Luo, C. and Gordon, V. D. 2016. Singly flagellated *Pseudomonas aeruginosa* chemotaxes efficiently by unbiased motor regulation. MBio 7:e00013.

58. Macnab, R. M. 1977. Bacterial flagella rotating in bundles: a study in helical geometry. Proc Natl Acad Sci U S A 74:221–5.

59. Vater, S. M., Weisse, S., Maleschlijski, S., Lotz, C., Koschitzki, F., Schwartz, T., Obst, U. and Rosenhahn, A. 2014. Swimming behavior of *Pseudomonas aeruginosa* studied by holographic 3D tracking. PLoS One 9:e87765.

60. Deziel, E., Comeau, Y. and Villemur, R. 2001. Initiation of biofilm formation by *Pseudomonas aeruginosa* 57RP correlates with emergence of hyperpiliated and highly adherent phenotypic variants deficient in swimming, swarming, and twitching motilities. J Bacteriol 183:1195–204.

61. Ha, D. G., Kuchma, S. L. and O’Toole, G. A. 2014. Plate-based assay for swimming motility in *Pseudomonas aeruginosa*. Methods Mol Biol 1149:59–65.

62. Martinez-Granero, F., Navazo, A., Barahona, E., Redondo-Nieto, M., Gonzalez de Heredia, E., Baena, I., Martin-Martin, I., Rivilla, R. and Martin, M. 2014. Identification of *flgZ* as a flagellar gene encoding a PilZ domain protein that regulates swimming motility and biofilm formation in *Pseudomonas*. PLoS One 9:e87608.

63. Petrova, O. E., Cherny, K. E. and Sauer, K. 2015. The diguanylate cyclase GcbA facilitates *Pseudomonas aeruginosa* biofilm dispersion by activating BdlA. J Bacteriol 197:174–87.

64. Huffer, S., Clark, M. E., Ning, J. C., Blanch, H. W. and Clark, D. S. 2011. Role of alcohols in growth, lipid composition, and membrane fluidity of yeasts, bacteria, and archaea. Appl Environ Microbiol 77:6400–8.

65. Cao, H., Wei, D., Yang, Y., Shang, Y., Li, G., Zhou, Y., Ma, Q. and Xu, Y. 2017. Systems-level understanding of ethanol-induced stresses and adaptation in E. coli. Sci Rep 7:44150.

66. Soufi, B., Krug, K., Harst, A. and Macek, B. 2015. Characterization of the E. coli proteome and its modifications during growth and ethanol stress. Front Microbiol 6:103.

67. Wolfe, A. J. and Berg, H. C. 1989. Migration of bacteria in semisolid agar. Proc Natl Acad Sci U S A 86:6973–7.

